# WWOX contributes to DNA damage, but not somatic instability in Huntington’s disease

**DOI:** 10.64898/2026.06.24.734331

**Authors:** Tiziana Petrozziello, Zachariah L. McLean, Adel Boudi, Sommer S. Huntress, Eric J. Granucci, Grace A. Field, Ranee Zara B. Monsanto, Ayleen L. Castillo Torres, Jennie C. L. Roy, Maheswaran Kesavan, Muzhou Wu, Neil Doherty, Ellen Sapp, Mahmoud A. Pouladi, Kimberly B. Kegel-Gleason, Marian DiFiglia, James F. Gusella, Ricardo Mouro Pinto, Ghazaleh Sadri-Vakili

**Author notes:** **Corresponding author:** Ghazaleh Sadri-Vakili, Ph.D., MassGeneral Institute for Neurodegenerative Disease, Massachusetts General Hospital, Bldg 114 16^th^ Street, R2200, Charlestown, MA 02129.

## Abstract

Huntington’s disease (HD), caused by a CAG repeat expansion in the huntingtin (*HTT*) gene, is characterized by progressive neurodegeneration and accumulation of DNA damage with multiple disease-modifier genes involved in DNA repair pathways. Previous studies have implicated ataxia telangiectasia mutated (ATM) signaling in the regulation of genomic stability and DNA damage repair (DDR) pathways in HD. ATM has also been linked to the WW domain-containing oxidoreductase (WWOX), a protein involved in DNA repair and maintenance of genomic stability, through the E3 ubiquitin ligase ITCH. However, whether this signaling pathway contributes to HD pathogenesis remains unknown. Here, we investigated the role of ATM-ITCH-WWOX signaling in HD. Our results revealed no significant alterations in total ATM, phosphorylated ATM (pATM-S1981), or ITCH in HD post-mortem prefrontal cortex (PFC) compared to controls. Although treatment of human neuroblastoma SH-SY5Y cells with HD PFC lysates did not alter pATM-S1981 levels, it increased histone H2AX phosphorylation at S139 (γ-H2AX), a marker of DNA double-strand breaks. This finding suggested the presence of persistent DNA damage signaling independent of canonical ATM activation. Conversely, WWOX levels were increased in both HD PFC and HD embryonic stem cell-derived cortical neurons. Additionally, treatment of SH-SY5Y cells with recombinant human WWOX protein or WWOX overexpression increased γ-H2AX levels, supporting a role for WWOX in promoting DNA damage. To determine whether WWOX contributed to DNA damage in HD, SH-SY5Y cells were treated with HD PFC lysates that were depleted of WWOX. Immuno-depletion of WWOX reduced the ability of HD PFC lysates to increase γ-H2AX, suggesting that WWOX contributes to DNA damage in HD. Finally, overexpression of WWOX in RPE1-AAVS1-CAG115 cells did not affect somatic CAG repeat instability, despite persistent increases in γ-H2AX levels. Collectively, our findings identify WWOX as a contributor to DNA damage in HD, acting independently of the ATM pathway.

## Introduction

Huntington’s disease (HD) is an autosomal hereditary disease caused by a CAG repeat expansion in exon 1 of *HTT*, resulting in an expanded polyglutamine (PolyQ) tract in HTT.^1,2^ CAG repeat length is a major determinant of HD onset and progression, with CAG repeats >36 considered to be pathogenic.^3^ CAG tract length is inversely correlated with age at disease onset and disease progression.^4–6^ Although CAG repeats in the normal range (<35 repeats) are stably transmitted, the CAG repeats in the mutant range are unstable and, when inherited, undergo contraction or expansion.^4–6^ This germline instability is mirrored by CAG length changes in somatic cells (somatic instability), a phenomenon modified by several genes, mainly involved in DNA repair.^6–18^ Although lowering HTT remains the primary therapeutic strategy, the discovery of HD modifiers has broadened the therapeutic landscape and multiple strategies targeting DNA repair modifier genes to delay disease onset and progression are currently under investigation.^13,17–19^

Ataxia telangiectasia mutated (ATM) signaling has also been implicated in the regulation of genomic instability and cellular response to DNA damage in HD.^20–22^ ATM is activated by DNA double-strand breaks and coordinates DNA repair, cell cycle regulation, and apoptosis.^23–26^ Previous studies demonstrated that mutant HTT can disrupt DNA repair through ATM signaling pathways, although the underlying mechanisms are not yet fully understood.^20–22^ Additionally, several proteins interacting with ATM have been implicated in the regulation of DNA damage repair (DDR) signaling: for example, it has been suggested that ATM functions together with the E3 ubiquitin ligase ITCH and the WW domain-containing oxidoreductase (WWOX) as part of a signaling axis involved in DNA repair.^27–29^

WWOX itself has been associated with transcriptional regulation,^30–33^ DNA repair,^27–29,34^ and maintenance of genomic stability,^28,34^ all key pathogenic processes disrupted in HD.^4–6^ Although WWOX was first described as a tumor suppressor gene,^35^ recent evidence highlighted its role in the development and diseases of the central nervous system (CNS).^36–43^ Mutations or altered expression of WWOX have been linked to several neurological and neurodegenerative disorders, including WWOX-related epileptic encephalopathy,^44–46^ autosomal recessive cerebellar ataxia 12,^47^ autism spectrum disorder, ^48,49^ Alzheimer’s disease (AD),^50^ multiple sclerosis (MS)^51^ and Parkinson’s disease (PD).^52^ Finally, WWOX was recently identified as a risk factor for PD progression^53^ and as a susceptibility locus for MS.^54^

Given the established role of DNA damage, DNA repair dysfunction, and somatic instability in HD pathogenesis, together with the proposed involvement of WWOX in ATM-associated DDR signaling, dysregulation of WWOX may represent a previously unexplored mechanism contributing to HD. Therefore, we investigated the involvement of ATM, ITCH, and WWOX in HD using post-mortem tissue and human cellular models to determine whether this pathway contributes to DNA damage and somatic instability in HD.

## Material and Methods

### Study approval

All methods were carried out in accordance with the guidelines and regulations of Mass General Brigham (MGB) and approved by the MGB licensing committees. Written informed consent was obtained from participants or the appropriate representative prior to tissue collection.

### Human tissue samples

Post-mortem prefrontal cortex (PFC) from control and HD patient brains were obtained by the Massachusetts Alzheimer’s Disease Research Center with approval from the MGB Institutional Review Board (IRB). In total, we assessed 13 control and 21 HD PFC. Demographic and genetic information, Vonsattel grade, and post-mortem interval (PMI) are summarized in Table 1.

**Table 1.** Post-mortem prefrontal cortex sample information.

|  | Sex | Age of death | PMI (h) | CAG repeat length |  | Proline |  | Vonsattel grade |
| --- | --- | --- | --- | --- | --- | --- | --- | --- |
|  |  |  |  | Allele 1 | Allele 2 | Allele 1 | Allele 2 |  |
| **Control 1** | F | 77 | 2 | Not determined | Not determined | Not determined | Not determined | Not determined |
| **Control 2** | M | 71 | 7 | Not determined | Not determined | Not determined | Not determined | Not determined |
| **Control 3** | F | 52 | 68 | Not determined | Not determined | Not determined | Not determined | Not determined |
| **Control 4** | M | 92 | 23 | Not determined | Not determined | Not determined | Not determined | Not determined |
| **Control 5** | M | 94 | Unknown | Not determined | Not determined | Not determined | Not determined | Not determined |
| **Control 6** | F | 58 | 18 | Not determined | Not determined | Not determined | Not determined | Not determined |
| **Control 7** | M | 60 | 14 | Not determined | Not determined | Not determined | Not determined | Not determined |
| **Control 8** | F | 89 | 8 | Not determined | Not determined | Not determined | Not determined | Not determined |
| **Control 9** | F | >90 | 24 | Not determined | Not determined | Not determined | Not determined | Not determined |
| **Control 10** | F | 77 | 72 | Not determined | Not determined | Not determined | Not determined | Not determined |
| **Control 11** | F | >90 | 24 | Not determined | Not determined | Not determined | Not determined | Not determined |
| **Control 12** | F | 79 | Unknown | Not determined | Not determined | Not determined | Not determined | Not determined |
| **Control 13** | F | >90 | 18 | Not determined | Not determined | Not determined | Not determined | Not determined |
| **HD 1** | M | 68 | 21.2 | 42 | 20 | 177 | 177 | 2 |
| **HD 2** | F | 51 | 19.67 | 44 | 19 | 177 | 177 | 3 |
| **HD 3** | M | 49 | 18.62 | 48 | 17 | 177 | 186 | 3 |
| **HD 4** | F | 40 | 20.5 | 50 | 20 | 177 | 177 | 3 |
| **HD 5** | F | 65 | 24.58 | 44 | 17 | 177 | 186 | 3 |
| **HD 6** | F | 56 | 21 | Not determined | Not determined | Not determined | Not determined | 4 |
| **HD 7** | F | 42 | 21.25 | 51 | 15 | 177 | 186 | 3 |
| **HD 8** | M | 74 | 9.11 | 41 | 23 | 177 | 177 | 3 |
| **HD 9** | M | 71 | 14.91 | 41 | 10 | 177 | 177 | 3 |
| **HD 10** | M | 51 | 23.42 | 49 | 19 | 177 | 177 | 3 |
| **HD 11** | M | 64 | 17.33 | 43 | 22 | 177 | 177 | 3 |
| **HD 12** | F | 53 | 17 | 42 | 22 | 177 | 177 | 3 |
| **HD 13** | M | 78 | 9.36 | 40 | 17 | 177 | 186 | 2 |
| **HD 14** | F | 65 | 9.5 | 43 | 17 | 177 | 186 | 4 |
| **HD 15** | M | 62 | 9 | 44 | 15 | 177 | 177 | 3 |
| **HD 16** | M | 48 | 16 | 49 | 18 | 177 | 186 | 4 |
| **HD 17** | F | 49 | 7.6 | 48 | 18 | 177 | 186 | 4 |
| **HD 18** | M | 36 | 18.63 | 51 | 15 | 177 | 186 | 3 |
| **HD 19** | F | 58 | Unknown | Not<br>determined | Not<br>determined | Not<br>determined | Not<br>determined | 4 |
| **HD 20** | F | 74 | 21 | Not<br>determined | Not<br>determined | Not<br>determined | Not<br>determined | 4 |
| **HD 21** | M | 41 | Unknown | Not<br>determined | Not<br>determined | Not<br>determined | Not<br>determined | 3 |

### Western blotting

Western blots were performed using previously described protocols.^55–57^ Briefly, 50μg of proteins was separated on a 4-20% Bis-Tris gel and transferred to a PVDF membrane using iBlot Dry Blotting System (Invitrogen, ThermoFisher, MA). Membranes were blocked with 5% milk in TBST and incubated overnight at 4°C with antibodies against ATM (1:500; #27156-1-AP, Proteintech, IL), ATM phosphorylated at S1981 (pATM-S1981; 1:500; #S1981, Cell Signaling, MA), ITCH (1:500; #12117S, Cell Signaling, MA), WWOX (1:500; #ab137726, Abcam, MA), and GAPDH (1:1 000; #MAB374, Millipore Sigma, Termecula, CA). Membranes were incubated with HRP-conjugated goat anti-rabbit or goat anti-mouse secondary antibodies (1:2 000; #111-035-003 and #115-035-166, Jackson ImmunoResearch Laboratories, West Grove, PA). Proteins were visualized using enhanced chemiluminescence (ThermoFisher Scientific, MA) and quantified using Image J (NIH, Bethesda, MD).

### Embryonic stem cells, and cortical neuron differentiations

Stem cell work was approved by the Human Embryonic Stem Cell Research Oversight (ESCRO) Committee and by the MGB Institutional Biosafety Committee (PIBC) (ESCRO #2015-01-02 and PIBC Reg #2017B000023). The isogenic human embryonic stem cells (ESCs) H9 (WA09, WiCell) - with a CAG repeat in the *HTT* alleles of 16/25 (encoding 18/27 polyglutamines) - were used to engineer new lines using Talen - containing 16/63 CAGs (encoding 18/65 polyglutamines).^58^ Human ESCs were grown on Matrigel (#354277, (Corning, MA) in mTeSR™ Plus (#100-0276, StemCell Technologies, Vancouver, Canada) and differentiated into cerebral cortex neurons using previously published protocols^59,60^ with minor modifications. Neural induction occurred on confluent ESCs plated on Matrigel (#354277, Corning, MA) for 10 days in Neural Induction Medium (NIM). Neuroepithelial sheets were lifted with dispase (#07923, StemCell Technologies, Vancouver, Canada), replated on Matrigel, and maintained in Neural Maintenance Medium (NMM) supplemented with 20ng/mL FGF (#PHG0023, Gibco, ThermoFisher Scientific, MA) until neural rosettes were observed. At day 42, the cells were cultured in NMM supplemented with 10ng/mL BDNF (#PHC7074, Gibco, ThermoFisher Scientific, MA) and 10ng/mL GDNF (#PHC7075, Gibco, ThermoFisher Scientific, MA) together with the anti-mitotic inhibitors 5-Fluoro-2’-Deoxyuridine (1µM; #F0503, Sigma-Aldrich, MO) and Uridine (1µM, #U3750, Sigma-Aldrich, MO).

### SH-SY5Y cells

Human neuroblastoma SH-SY5Y cells (ATCC® CRL-2266) were cultured in DMEM/Nutrient Mixture F-12 supplemented with 10% inactivated FBS, 2mM L-glutamine, 50μg/mL streptomycin, and 50IU/mL penicillin (#A5670701, Gibco, ThermoFisher Scientific, MA). The cells were kept at 37°C and 5% CO_2_

### SH-SY5Y cell culture treatments

#### HD PFC treatment

Cells were treated with 10 or 25ng/mL lysates derived from control (n=3) or HD PFC (n=3) for 24h before assessing histone H2AX phosphorylated at S139 (γ-H2AX) levels by immunocytochemistry/immunofluorescence (ICC/IF).

#### Recombinant protein treatment

Recombinant full-length human WWOX protein (rWWOX) was purchased from Pepmic Co (Suzhou, China). Cells were incubated with 1, 10, 100 or 500ng/mL rWWOX for 24h or 48h before measuring cell viability by MTT assay or with 10, 100 or 150ng/mL for 24h before assessing γ-H2AX levels by ICC/IF.

#### SH-SY5Y overexpressing WWOX cells

PiggyBac plasmids for WWOX overexpression and control were generated by VectorBuilder: pPB[Exp]-mCherry/Neo-EF1A>hWWOX[NM_016373.4]/FLAG (VB240621-1305qkd) and pPB[Exp]-mCherry/Neo-EF1A>ORF_Stuffer (VB900167-6129xyz). Detailed vector information can be accessed using these IDs at vectorbuilder.com. Cells were transfected with 200ng/mL plasmid in lipofectamine 2000 (ThermoFisher, MA) using previously published protocols.^56,61^ Stable cells were selected with 500ng/mL G418 (#AAJ63871AB, Fisher Scientific, MA), and clone selection was monitored visualizing mCherry fluorescent signal at a Zoe fluorescent cell imager (Biorad Laboratories, MA). Efficiency of transfection was determined by western blotting.

#### Small interfering RNA (siRNA)

WWOX knockdown was performed with siGENOME Human WWOX siRNA (siWWOX) (#D-003961-15-0005, Horizon Discovery, GE Healthcare Dharmacon, CO), using a siGENOME non-targeting control pool (#D-001206-13-05, Horizon Discovery, GE Healthcare Dharmacon, CO) as control. SH-SY5Y cells were transfected with 10nM siWWOX in Lipofectamine 2000 (ThermoFisher, MA) for 48h, as previously described.^56^ Knockdown efficiency was confirmed by western blotting.

#### 4,5-dimethylthiazol-2-yl]-2,5-diphenyl-tetrazolium bromide (MTT) assay

SH-SY5Y cells were incubated with MTT for 1h at 37°C and collected in dimethyl-sulfoxide (DMSO). Optical density was measured at 620nm by using a Microplate reader (Synergy 2, Biotek Instruments).^61^

#### ICC/IF and image analysis

Cells were plated on Millicell EZ SLIDE glass slides (Millipore Sigma, CA), treated as described above, and fixed with 4% paraformaldehyde (PFA) using previously published protocols.^56^ Next, cells were permeabilized with 0.15% Triton-X in PBS and blocked in 0.5% BSA in PBS-Tween 20 (PBS-T). Cells were incubated overnight with pATM-S1981 (1:200; Cell Signaling, MA) or γ-H2AX (1:1 000; Cell Signaling, MA) antibodies followed by incubation in Alexa Flour-conjugated secondary antibodies (goat-anti-rabbit A568, donkey-anti-mouse 488, goat-anti-rabbit 488; 1:1 000; ThermoFisher Scientific, MA) and DAPI. Slides were coverslipped with Vectashield Antifade Mounting Medium (Vector Labs, CA). Z-stack were acquired using a Zeiss LSM 880 confocal microscope (Zeiss Microscope, MA) with a 40x objective. At least 4 random images were captured per condition, and analyzed in ImageJ (NIH, Bethesda, MD).

#### WWOX immuno-depletion

WWOX content was immuno-depleted from three HD PFC homogenates with the highest WWOX levels, as determined by western blotting, using a previously published protocol.^62^ Briefly, 150μg of proteins from each HD PFC was incubated with anti-WWOX antibody (#ab137726, Abcam, MA) and magnetic beads (Invitrogen, ThermoFisher, MA). Supernatants, depleted of WWOX, were collected using a magnetic rack and saved as HD-, whereas bead-bound WWOX-enriched fractions were saved d as HD+. Immuno-depletion efficiency was determined by western blotting.

#### RPE-1AAVS1-CAG115 overexpressing WWOX cells

PiggyBac plasmids for WWOX overexpression and control were generated as described above. Briefly, RPE-1 AAVS1-CAG115 cells^15^ were transfected by nucleofection with WWOX overexpression or control plasmids together with Super PiggyBac Transposase (System Biosciences, CA). Nucleofection was performed using the 4D-Nucleofector X Unit (Lonza, MA) with P3 4D-Nucleofector X Solution (V4XP-3024) and the EA-104 program. Following transfection, cells were divided into six parallel populations and selected with 500μg/mL G418 (#10131035, Gibco, ThermoFisher, MA).

#### Measurement of repeat instability

RPE-1 AAVS1-CAG115 cells were plated in 96-well plates at 50,000 cells/well under confluent, non-proliferative conditions. Baseline samples were collected at day 0, and two replicate wells per population were cultured for 14, 28 or 42 days. Genomic DNA was extracted using the Quick-DNA 96 Kit (Zymo Research, CA). The repeat region was amplified using a nested PCR strategy designed to selectively amplify the transgene. In the first PCR, a region spanning HTT to GFP was amplified using the Taq PCR Core Kit with Q-Solution (Qiagen, MD) and primers ATGAAGGCCTTCGAGTCCCTCAAGTCCTTC (forward) and GTCCAGCTCGACCAGGATG (reverse). Reactions were performed using 5μL genomic DNA under the following conditions: 95°C for 5min; 12 cycles of 95°C for 30s, 65°C for 30s, and 72°C for 1min 30s; followed by a final extension at 72°C for 10min. For repeat sizing, a second PCR was carried out using 2μL of the first-round product as template, with primers /56-FAM/ATGAAGGCCTTCGAGTCCCTCAAGTCCTTC (forward) and GGCGGCTGAGGAAGCTGAGGA (reverse) for 22 cycles. PCR products were analyzed by capillary electrophoresis on a 3730XL DNA Analyzer. Fragment analysis data were process using the TRACE R package to generate somatic instability metrics.^15^ We used a peak threshold of 5% and limited the window of analysis to 15 repeats around the index peak (starting repeat length).

#### Statistics

Statistical analyses were performed using GraphPad Prism. Data are presented as individual plots with median and interquartile range or as mean ± standard error of the mean (SEM), as indicated in the figure legends. Comparison between two groups were performed using Mann-Whitney U test. Comparisons among multiple groups were performed using one-way or two-way ANOVA followed by Tukey’s multiple comparisons test. Correlations between WWOX levels and age at death or CAG repeat were performed using Spearman correlations. To assess the relationship between repeat change, time in culture (div), and WWOX overexpression (group), multiple linear regression models were fit in R using the formula average_repeat_change∼div+group. All tests were two-sided, and statistical significance was set at p<0.05. Exact p values are reported throughout.

## Results

### ATM signaling is not altered in HD despite increases in γ-H2AX levels

To determine whether ATM signaling is altered in HD, we first assessed the levels of ATM, its activated form (pATM-S1981), and ITCH, a downstream effector of ATM,^63^ in human post-mortem HD and control PFC samples by western blotting. There were no differences in total ATM (Figure 1A), pATM-S1981 (Figure 1B), or ITCH levels (Figure 1C) between HD and control PFC.

**Figure 1.**
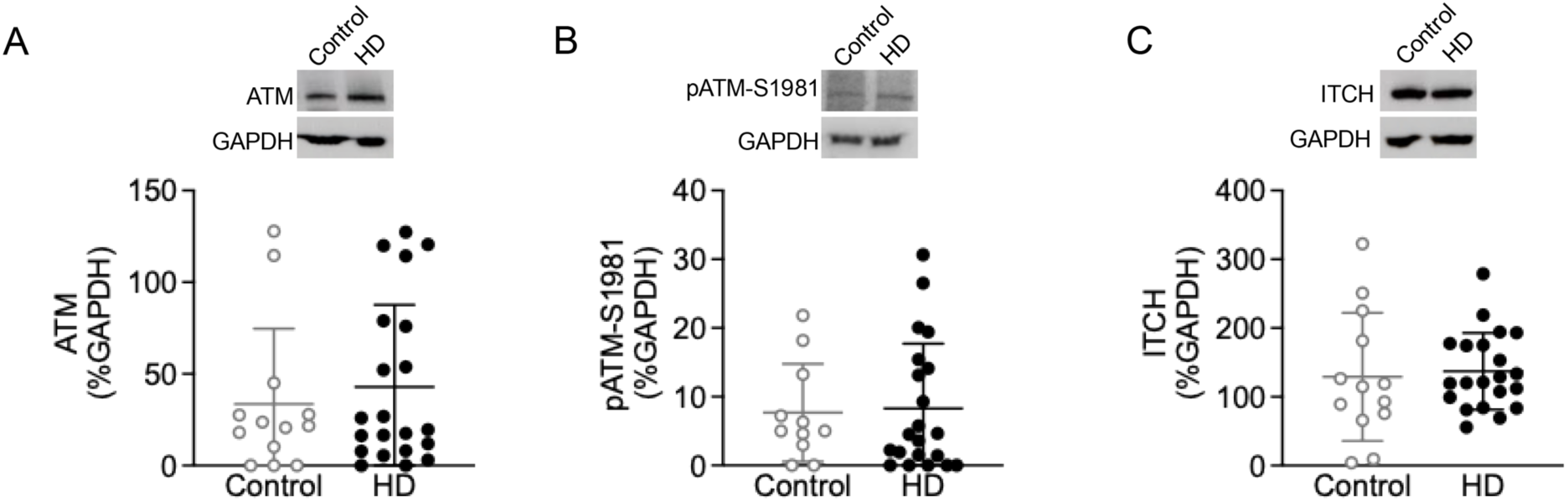
ATM, pATM-S1981, and ITCH levels were unchanged in HD post-mortem PFC. **(A)** ATM (Mann-Whitney U test=130, p=0.8337), **(B)** pATM-S1981 (Mann-Whitney U test=105, p=0.6944), and **(C)** ITCH (Mann-Whitney U test=118, p=0.5288) levels were unchanged in HD PFC (n=21) compared to control PFC (n=13). Data are shown as mean±SEM. p>0.05.

Next, we assessed pATM-S1981 and its downstream target, γ-H2AX,^64,65^ a marker of DNA double-strand breaks^66^ in SH-SY5Y cells treated with HD PFC lysates by ICC/IF. Camptothecin (CPT; 10μM/2h), a DDR activator,^67,68^ served as a positive control.

Consistent with our findings in HD PFC, treatment with HD lysates (25ng/mL/24h) did not alter pATM-S1981 levels in SH-SY5Y cells (Figure 2A and B). In contrast, γ-H2AX levels were increased following HD treatment (Figure 2A and C). Additionally, γ-H2AX did not colocalize with pATM-S1981 (Figure 2D-G), suggesting that HD-induced γ-H2AX accumulation occurs in the absence of alterations in ATM signaling.

**Figure 2.**
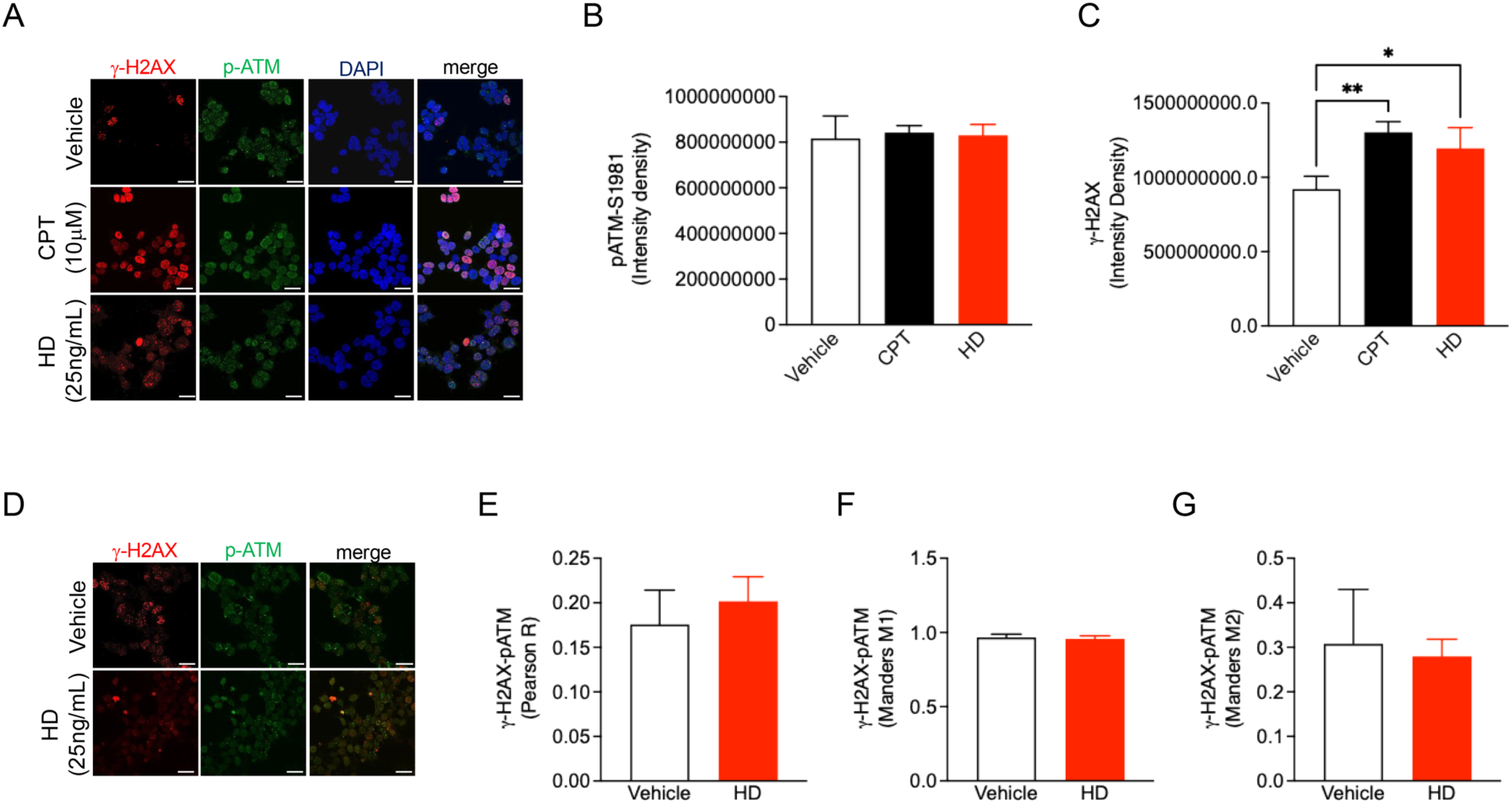
Treatment with HD PFC lysates increasedγ-H2AX levels but not pATM-S1981 levels in SH-SY5Y cells. **(A)** Representative immunofluorescence images of γ-H2AX (red), pATM-S1981 (green), DAPI (blue), and merge (right) in SH-SY5Y cells following treatment with HD PFC lysates (25ng/mL/24h). **(B)** One-way ANOVA revealed no significant effect of treatment on pATM-S1981 levels [F(2,6)=0.1127; p=0.8953]. Neither CPT treatment (10μM/2h) nor HD lysates altered pATM-S1981 levels compared to vehicle-treated cells (n=3 biological replicates; Tukey’s test, p=0.8791, and p=0.9614, respectively). **(C)** One-way ANOVA revealed a significant effect of treatment on γ-H2AX levels [F(2,6)=10.87; p=0.0101]. Both CPT treatment (10μM/2h) and HD lysates significantly increased γ-H2AX levels compared to vehicle-treated cells (n=3 biological replicates; Tukey’s test, p=0.0080, and p=0.0347, respectively). **(D)** Representative immunofluorescence images of γ-H2AX (red), pATM-S1981 (green), and merge (right) in SH-SY5Y following treatment with HD PFC lysates (25ng/mL/24 h). HD lysates did not alter γ-H2AX and pATM-S1981 co-localization in SH-SY5Y cells as assessed by **(E)** Pearson’s correlation coefficient (Mann-Whitney U test=2.500, p=0.5000), **(F)** Manders’ M1 coefficient (Mann-Whitney U test=4, p>0.9999), and **(G)** Manders’ M2 coefficient (Mann-Whitney U test=3, p=0.7000). Data are shown as mean±SEM. Scale bar 50μm. * p<0.05; ** p<0.01.

To determine whether γ-H2AX increase was specific to HD lysates and not driven by exposure to human brain material or toxic protein concentration, cells were treated with either a lower concentration of HD PFC lysates (10ng/mL/24h) or control PFC lysates (25ng/mL/24h). While γ-H2AX levels were increased following CPT treatment and treatment with HD lysates (25ng/mL/24h), neither control PFC lysates nor the lower concentration of HD lysates altered γ-H2AX levels compared to vehicle-treated cells (Supplementary Figure 1), indicating that the increase in γ-H2AX is both concentration-dependent and specific to HD.

### WWOX levels are increased in human post-mortem HD PFC but do not correlate with CAG repeat expansion

Given the role of WWOX in ATM signaling and DNA damage pathways,^27–29,34^ we next assessed whether its levels were altered in HD using western blots. WWOX levels were increased in HD PFC compared to control PFC (Figure 3A). However, WWOX levels did not correlate with either CAG repeat length (Figure 3B) or age at death (Figure 3C) in HD samples.

**Figure 3.**
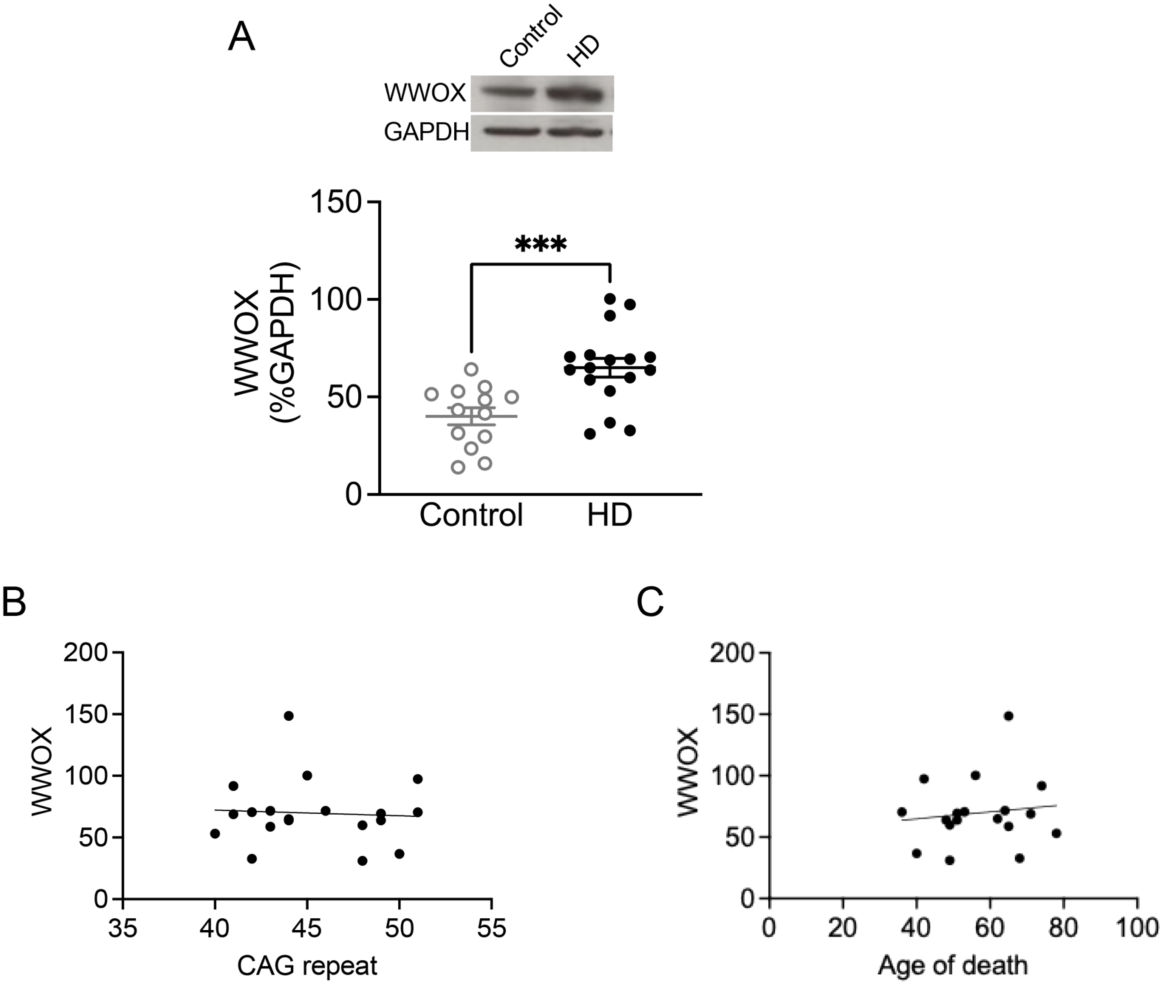
WWOX levels were increased in human HD post-mortem PFC but did not correlate with CAG repeat length or age at death. **(A)** WWOX levels were significantly increased in HD PFC (n=18) compared to control PFC (n=13) (Mann-Whitney U test=31, p=0.0005). **(B)** WWOX levels did not correlate with CAG repeat length in PFC (Spearman’s correlation, r=0.05302, p=0.8397). **(C)** WWOX levels did not correlate with age at death in HD PFC (Spearman’s correlation, r=0.05891, p=0.8164). Data are shown as mean±SEM. *** p<0.001.

### Increases in WWOX induce neuronal toxicity in HD ESC-derived cortical neurons

We assessed WWOX levels in cortical neurons derived from HD ESCs and isogenic controls. WWOX levels were increased in HD ESC-derived cortical neurons (65Q) compared to controls (27Q) (Figure 4A).

**Figure 4.**
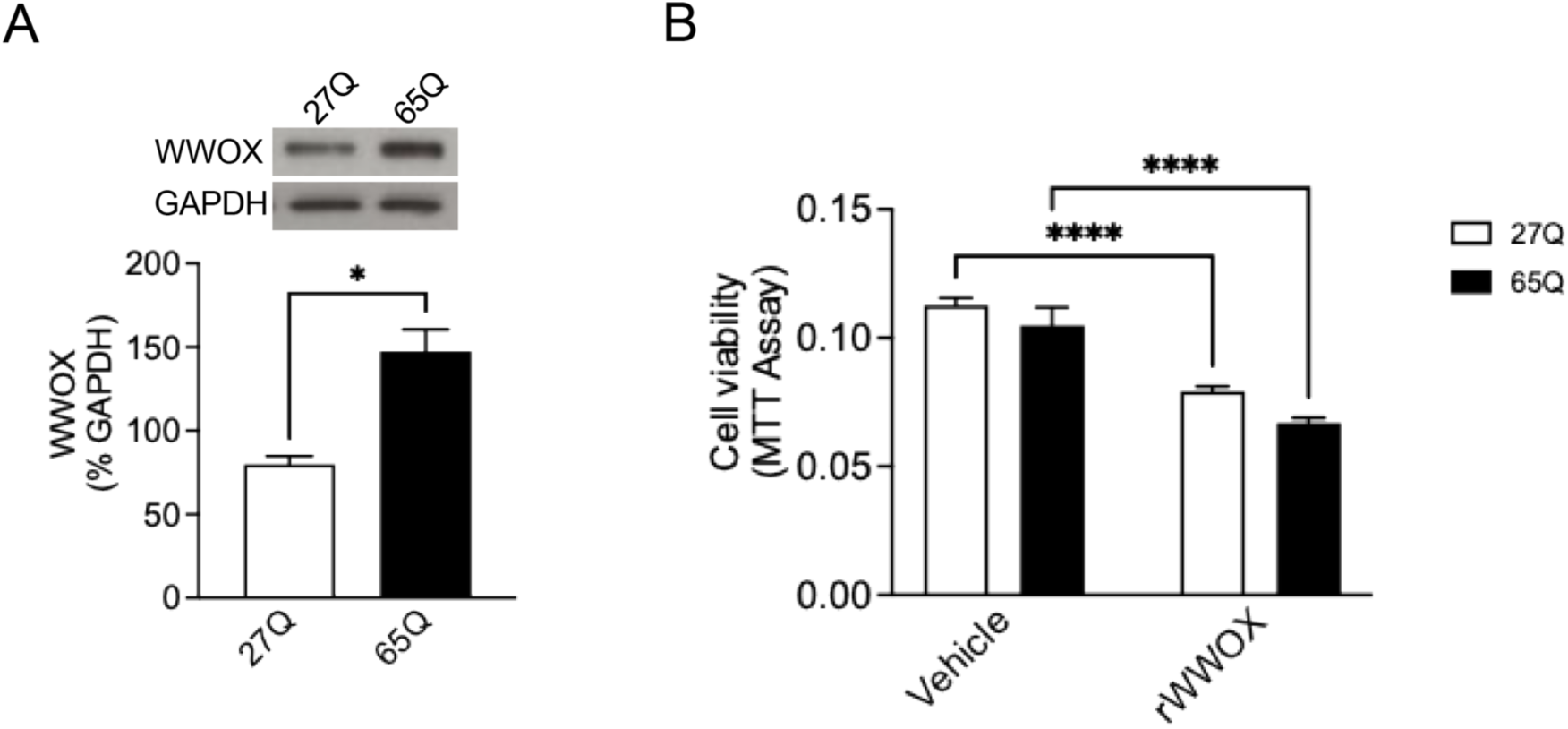
WWOX levels were increased in human HD ESC-derived cortical neurons, and rWWOX induced to cell death. **(A)** WWOX levels were significantly increased in HD ESC-derived cortical neurons (65Q) compared to isogenic control lines (27Q) (n=4 biological replicates; Mann-Whitney U test=0, p=0.0266). **(B)** Treatment with rWWOX (100ng/mL) induced cell death in both HD ESC-derived cortical neuron (65Q) and isogenic control neurons (27Q). Two-way ANOVA revealed a significant effect of treatment [F(1, 28)=78.44; p<0.0001], and genotype [F(1,28)=6.181; p=0.0192], with no significant treatment x genotype interaction [F(1,28)=0.3129; p=0.5804]. Specifically, treatment with 100 ng/mL of rWWOX significantly reduced cell viability in both HD ESC-derived cortical neurons (65Q) (Tukey’s test, p<0.0001) and isogenic control cells (27Q) (Tukey’s test, p<0.0001) compared to their respective vehicle-treated controls. Data are shown as mean±SEM. * p<0.05; **** p<0.0001.

To determine the consequences of increased WWOX, SH-SY5Y cells were treated with increasing concentrations of a human rWWOX protein (1, 10, 100 and 500ng/mL) for 24h and 48h prior to measuring cell viability using the MTT assay.

Treatment with rWWOX caused cell death in a dose- and time-dependent manner in SH-SY5Y cells, with significant decreases observed at 100 or 500ng/mL after 24h and at 10, 100 or 500ng/mL after 48h (Supplementary Figure 2). Based on these results, HD and control ESC-derived cortical neurons were treated with 100ng/mL rWWOX for 24h. Our results revealed a significant decrease in cell viability in both rWWOX-treated HD cortical neurons (65Q) and their isogenic controls (27Q) compared to the respective vehicle-treated cells (Figure 4B).

### Elevated WWOX increases γ-H2AX levels in SH-SY5Y cells

To determine whether WWOX contributes to DNA damage, we measured γ-H2AX in SH-SY5Y cells following treatment with rWWOX (10, 100 or 150ng/mL) by ICC/IF using CPT as a positive control. γ-H2AX levels were increased in CPT-treated cells and in cells treated with either 100 or 150ng/mL of rWWOX compared to vehicle-treated cells (Figure 5A and B).

**Figure 5.**
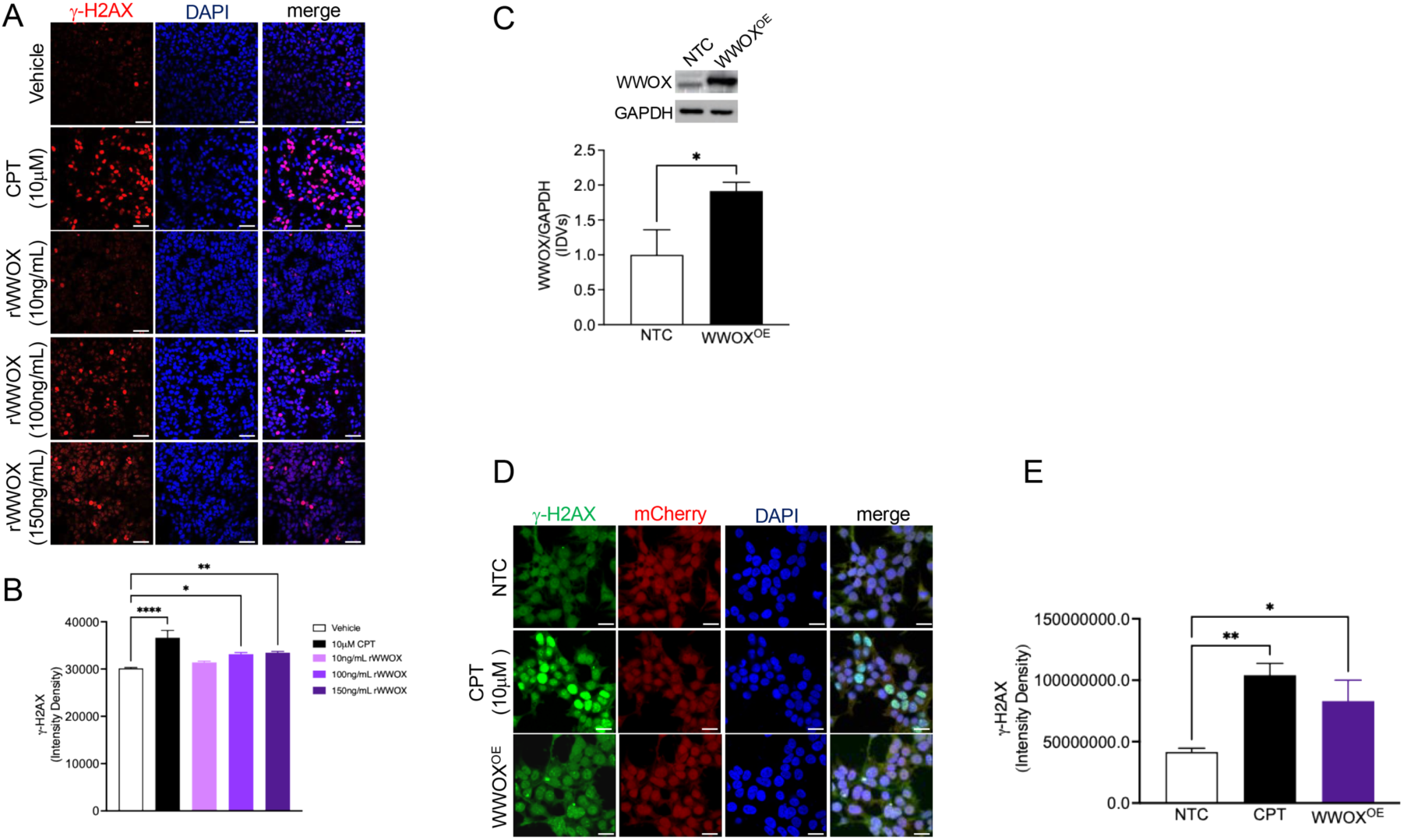
Elevated WWOX increased γ-H2AX levels in SH-SY5Y cells. **(A)** Representative immunofluorescence images of γ-H2AX (red), DAPI (blue), and merge (right) in SH-SY5Y cells following treatment with rWWOX. **(B)** One-way ANOVA revealed a significant effect of treatment on γ-H2AX [F(4,42)=13.96; p<0.0001]. CPT treatment (10μM/2h) significantly increased γ-H2AX levels in SH-SY5Y cells (Tukey’s test, p<0.0001). While treatment with 10ng/mL of rWWOX did not alter γ-H2AX levels (Tukey’s test, p=0.6018), treatment with either 100ng/mL (Tukey’s test, p=0.0139) or 150ng/mL of rWWOX (Tukey’s test, p=0.0038) significantly increased γ-H2AX levels compared to vehicle-treated cells. **(C)** WWOX levels were significantly increased in SH-SY5Y overexpressing WWOX (WWOX^OE^) cells compared to NTC (Mann-Whitney U test=1, p=0.0476). **(D)** Representative immunofluorescence images of γ-H2AX (green), mCherry (red), DAPI (blue), and merge (right) in SH-SY5Y WWOX^OE^ cells and NTC. **(E)** One-way ANOVA revealed a significant effect of treatment on γ-H2AX levels [F(2,12)=8.643; p=0.0047]. CPT treatment (10μM/2h) significantly increased γ-H2AX levels in SH-SY5Y cells (Tukey’s test, p=0.0037). Similarly, γ-H2AX levels were significantly increased in SH-SY5Y WWOX^OE^ cells compared to NTC (Tukey’s test, p=0.0314). Data are shown as mean±SEM. Scale bar 50μm. * p<0.05; ** p<0.01; **** p<0.0001.

Next, we generated SH-SY5Y cells stably overexpressing WWOX (WWOX^OE^). WWOX overexpression was confirmed by western blotting demonstrating higher WWOX levels in WWOX^OE^ compared to non-transfected control (NTC) cells (Figure 5C). To further determine the effects of WWOX overexpression on DNA damage, we measured γ-H2AX levels by ICC/IF. γ-H2AX levels were increased in both CPT-treated cells and WWOX^OE^ cells compared to NTC cells (Figure 5D and E).

To determine whether reducing WWOX levels could mitigate γ-H2AX accumulation, WWOX levels were knocked down using siWWOX prior to measuring γ-H2AX levels by ICC/IF. WWOX knockdown was confirmed by western blotting demonstrating lower WWOX levels in siWWOX-treated cells compared to siControl-treated cells (Supplementary Figure 3A). While CPT treatment increased γ-H2AX levels, γ-H2AX levels were decreased in siWWOX-treated cells compared to both siControl- and CPT-treated cells (Supplementary Figure 3B and C).

### WWOX immuno-depletion reduces HD-induced γ-H2AX accumulation

To determine whether endogenous WWOX levels from HD PFC lysates directly contribute to DNA damage, WWOX was immuno-depleted from the samples using an anti-WWOX antibody. Immuno-depletion was confirmed by western blotting (Figure 6A). The results demonstrated a significant decrease in WWOX levels in WWOX-depleted lysates (HD-), compared to HD and WWOX-enriched (HD+) lysates (Figure 6B).

**Figure 6.**
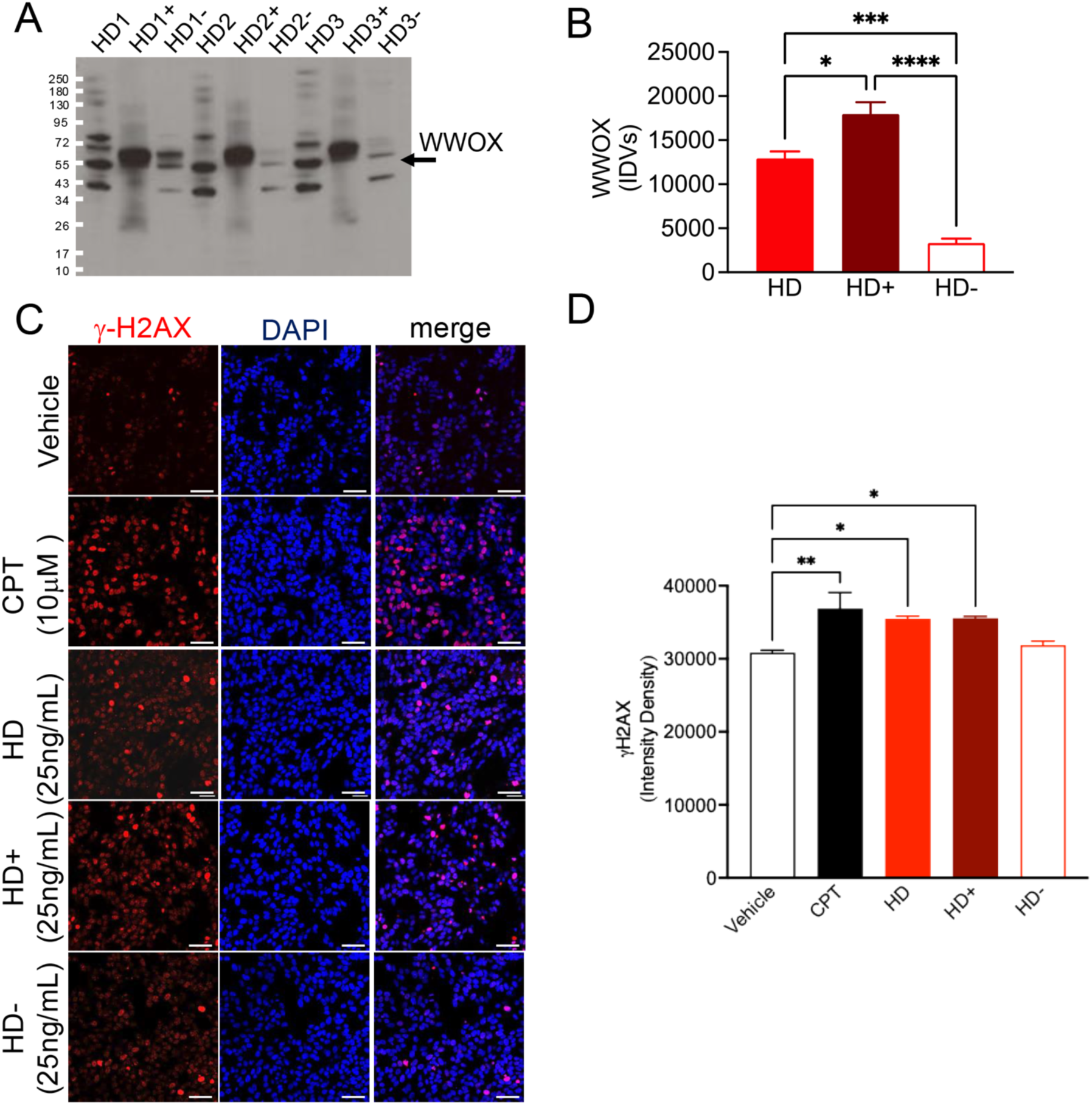
WWOX immuno-depletion reduced HD-induced γ-H2AX accumulation in SH-SY5Y cells. **(A)** Representative western blot image of WWOX levels following immuno-depletion in human post-mortem HD PFC (n=3). **(B)** One-way ANOVA revealed a significant effect of immuno-depletion on WWOX levels [F(4,6)=62.79; p<0.0001]. WWOX levels were significantly increased in HD+ PFC compared to HD PFC (Tukey’s test, p=0.0210). In contrast, WWOX levels were significantly decreased in WWOX-depleted (HD-) PFC compared to both HD PFC (Tukey’s test, p=0.0009) and WWOX-enriched (HD+) PFC (Tukey’s test, p<0.0001). **(C)** Representative immunofluorescence images of γ-H2AX (red), DAPI (blue), and merge (right) in SH-SY5Y cells following treatment with HD, HD+, or HD- PFC lysates. **(D)** One-way ANOVA revealed a significant effect of treatment on γ-H2AX levels [F(4,27)=5.699; p=0.0019]. CPT treatment (10μM/2h) significantly increased γ-H2AX levels compared to vehicle-treated cells (Tukey’s test, p=0.0037). Similarly, treatment with either HD PFC lysates (Tukey’s test, p=0.0473) or HD+ PFC lysates (Tukey’s test, p=0.0424) significantly increased γ-H2AX levels compared to vehicle treated cells. In contrast, treatment with HD- PFC lysates did not alter γ-H2AX levels compared to vehicle treated cells (Tukey’s test, p=0.9608). Data are shown as mean±SEM. Scale bar 50μm. * p<0.05; ** p<0.01; *** p<0.001; **** p<0.0001.

Next, we measured γ-H2AX levels by ICC/IF in SH-SY5Y cells following treatment with HD, HD+, and HD- lysates. γ-H2AX levels were increased in CPT-treated cells compared to vehicle-treated cells. Treatment with HD or HD+ lysates (25ng/mL/24h) increased γ-H2AX levels, whereas HD- lysates had no effect (Figure 6C and D).

### WWOX overexpression increases γ-H2AX levels but does not alter repeat instability in RPE-1AAVS1-CAG115 cells

To determine whether WWOX contributed to CAG repeat instability in HD, we used a newly-established cellular model, the RPE-1AAVS1-CAG115 cells.^15^

Six populations of RPE-1AAVS1-CAG115 cells stably overexpressing WWOX were generated, and WWOX overexpression was confirmed by western blotting revealing higher WWOX levels in all six clones compared to NTC cells (Figure 7A).

**Figure 7.**
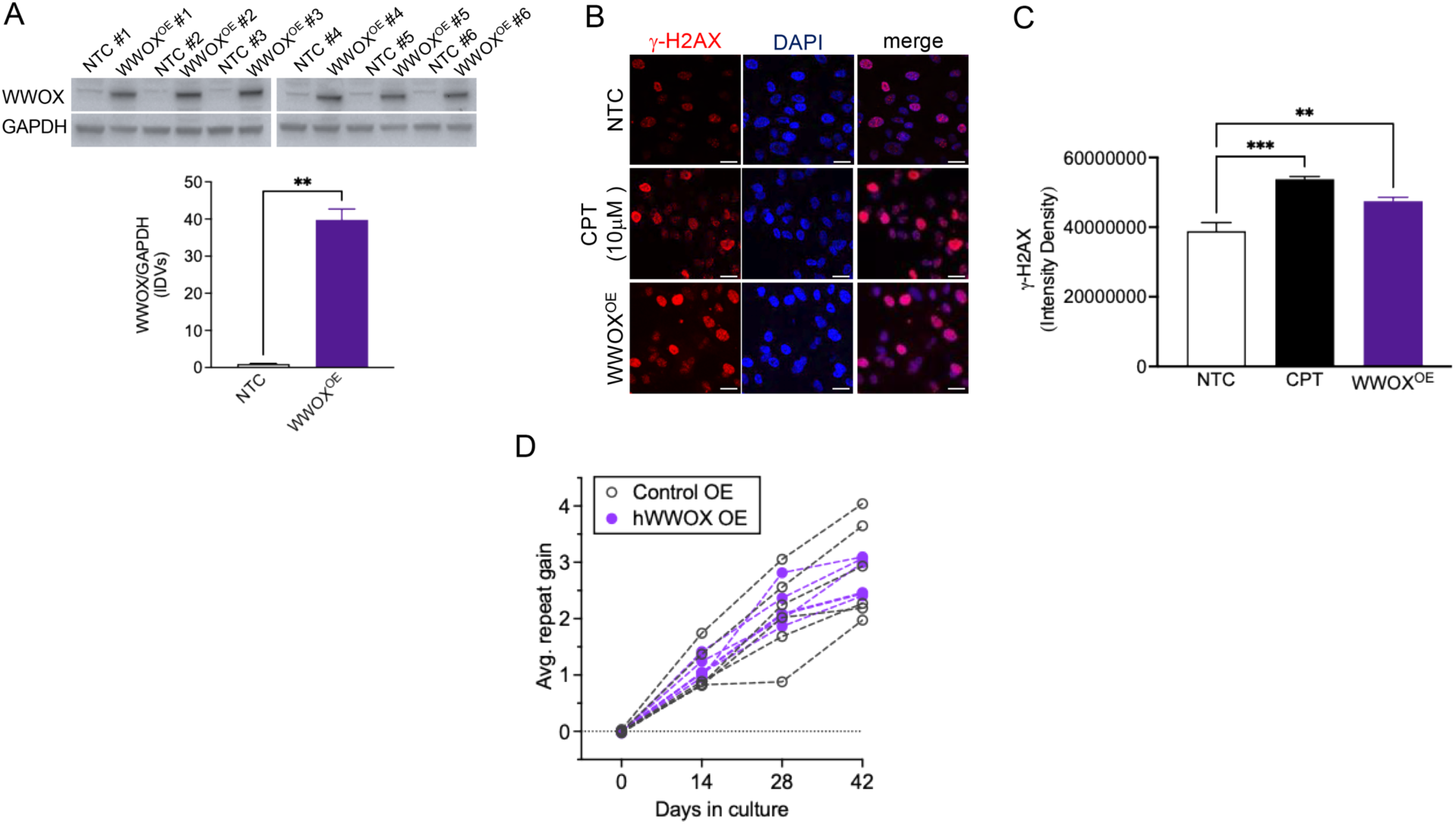
WWOX overexpression increased γ-H2AX levels but did not alter CAG repeat length in RPE-1AAVS1-CAG115 cells. **(A)** Representative western blot image of WWOX levels in six clones of RPE-1AAVS1-CAG115 cells. WWOX levels were significantly increased in RPE-1AAVS1-CAG115 stably overexpressing WWOX cells (WWOX^OE^) compared to non-transfected control (NTC) (Mann-Whitney U test=0, p=0.0022). **(B)** Representative immunofluorescence images of γ-H2AX (red), DAPI (blue), and merge (right) in in RPE-1AAVS1-CAG115 cells overexpressing WWOX (WWOX^OE^). **(C)** One-way ANOVA revealed a significant effect of treatment on γ-H2AX levels [F(2,11)=18.07; p=0.003]. CPT treatment (10μM/2h) significantly increased γ-H2AX levels in RPE-1AAVS1-CAG115 cells compared to NTC cells (Tukey’s test, p=0.0002). Similarly, γ-H2AX levels were significantly increased in RPE-1AAVS1-CAG115 cells overexpressing WWOX (WWOX^OE^) compared to NTC (Tukey’s test, p=0.0081). **(D)** Representative traces of average repeat gain measured over time in RPE-1AAVS1-CAG115 cells overexpressing WWOX (WWOX^OE^). Multiple linear regression was used to assess repeat change. Days in vitro showed a significant effect (p<2×10⁻¹⁶), whereas WWOX overexpression had no effect (p=0.877). Data are shown as mean±SEM. Scale bar 50μm. ** p<0.01; *** p<0.001.

Next, we assessed γ-H2AX levels in all cell lines using ICC/IF. γ-H2AX levels were increased in RPE-1AAVS1-CAG115 cells overexpressing WWOX (WWOX^OE^) compared to NTC cells, similar to CPT-treated cells (Figure 7B and C).

Lastly, repeat instability was measured after 0, 14, 28 or 42 days in culture in all populations. WWOX overexpression did not alter the rate of CAG repeat instability at any time point (Figure 7D).

## Discussion

In this study, we investigated the ATM-ITCH-WWOX signaling axis and identified WWOX as a contributor to DNA damage in HD. While ATM, pATM-S1981, and ITCH levels were unchanged, WWOX levels were increased in HD PFC and HD ESC-derived cortical neurons. Elevated WWOX consistently increased γ-H2AX levels across multiple cellular models, but did not alter CAG repeat instability, suggesting that WWOX contributes to DNA damage but not somatic instability in HD.

DNA damage and alterations in DNA repair are increasingly recognized as key features of HD pathogenesis.^4–6^ Genome-wide association studies have indeed identified several disease modifier genes that are involved in DNA repair mechanisms, in particular DNA mismatch repair (MMR), including *MSH3*, *MLH1*, *PMS1*, *PMS2*, *LIG1*, and *FAN1* to name a few.^6–18^ However, alterations in the DNA repair machinery were linked to HD pathogenesis well before the discovery of the MMR disease modifier genes. Mutant HTT itself has been shown to disrupt DNA repair mechanisms,^20–22,69,70^ including ATM signaling.^20–22^ ATM is activated upon DNA damage^65,71^ or oxidative stress^72^ and promotes phosphorylation of histone H2AX at sites of DNA damage.^64,65^ However, despite previous reports linking mutant HTT to ATM signaling, we found no alterations in ATM and pATM-S1981 levels, nor in the levels of their downstream effector ITCH^63^ in HD post-mortem PFC. Similarly, treatment with HD brain lysates increased γ-H2AX levels without affecting pATM-S1981 levels in SH-SY5Y cells. Together, these findings suggest that increased DNA damage signaling occurs independently of ATM activation. This discrepancy between our findings and the previously published results could be due to differences in the biological systems examined across studies and may indicate that ATM dysregulation is not consistent across HD models or disease stage.

An interesting component of ATM signaling is represented by WWOX, whose role in HD has remained unexplored. Upon activation, WWOX regulates ATM-mediated DDR in a feed-forward loop.^27–29^ Specifically, loss of WWOX delays ATM activation and impairs DNA repair. Surprisingly, our findings demonstrate that WWOX is increased in HD post-mortem brain and human cellular models. Because the post-mortem cohort consisted primarily of Vonsattel grade 3-4 cases, WWOX upregulation may be associated with late-stage HD pathology. Elevated WWOX levels did not correlate with age at death or CAG repeat length, indicating that WWOX upregulation is not simply driven by aging or repeat expansion. In contrast to reports describing reduced WWOX expression in AD,^50^ and MS,^51^ WWOX was increased in HD, suggesting disease-specific dysregulation. Whether increased WWOX represents a compensatory response to chronic genomic stress or a maladaptive process contributing to neuronal dysfunction in HD remains to be determined.

Multiple complementary approaches used in this study supported a direct role for WWOX in DNA damage signaling in HD. rWWOX treatment and stable WWOX overexpression increased γ-H2AX levels, whereas siWWOX and WWOX immunodepletion reduced HD-associated γ-H2AX accumulation. These observations support a mechanistic role for elevated WWOX in promoting DNA damage signaling. Although previous studies have linked WWOX deficiency to genomic instability and impaired DDR signaling,^27–29,34^ our findings indicate that elevated WWOX can also promote DNA damage. Together, these observations suggest that maintaining appropriate WWOX levels may be important for cellular homeostasis.

Another intriguing aspect of this study is that WWOX-induced DNA damage signaling did not translate into changes in somatic instability. Although WWOX overexpression consistently increased γ-H2AX levels, it did not alter CAG repeat length dynamics in RPE-1AAVS1-CAG115 cells. This is consistent with our findings in post-mortem HD PFC, where WWOX levels did not correlate with CAG repeat length. Our findings suggest that elevated WWOX may influence HD pathogenesis through cellular stress responses, genomic damage signaling, or neuronal vulnerability rather than direct modulation of CAG repeat expansion.

Several limitations should be acknowledged. The post-mortem cohort consisted primarily of individuals with advanced neuropathology, limiting our ability to determine when WWOX dysregulation emerges during disease progression. In addition, γ-H2AX is a well-established marker of DNA damage signaling but does not directly quantify DNA double-strand breaks. The relationship between WWOX dysregulation and mutant HTT also remains unclear. Whether WWOX acts downstream of mutant HTT, contributes independently to disease pathogenesis, or represents a broader response to neurodegeneration remains to be determined. Defining the relationship between mutant HTT and WWOX will be important to determine whether modulation of WWOX can mitigate disease-relevant phenotypes.

Collectively, our findings identify WWOX as a novel contributor to DNA damage signaling in HD despite having no detectable effect on somatic instability. These results expand our understanding of DNA damage pathways in HD, support a previously unrecognized role for WWOX in HD-associated DNA damage, and warrant further investigation into the mechanisms linking WWOX dysregulation to HD pathogenesis.

## Availability of Data and Materials

The data supporting the findings of this study are available within the article and/or its supplementary material.

## Acknowledgements

The authors would like to thank patients and their families for sample donation. MAP was supported by the BC Children’s Health Research Institute Investigator Grant Award (IGAP), and grants from the Canadian Institutes of Health Research (CIHR grant # ENG-191555) and Huntington’s Disease Foundation (HDF grant # 1410890). JFS was supported by NINDS (NS091161). RMP was supported by the National Institute of Health grant NS126420. RMP and GSV were supported by the Dake Family Foundation.

## Author contributions

TP contributed to study design, data collection, data analysis and drafting of the manuscript. ZLM, AB, SSH, EJG, GAF, RZBM, ALCT, JCLR, MK, MW, ND, and ES contributed to data collection, data analysis, and editing of the manuscript. MAP, KKG, MDF, JFG, and RMP contributed to study design and editing of the manuscript. GSV contributed to study design, supervised the study and drafted the manuscript.

---

**Supplementary Figure 1.**
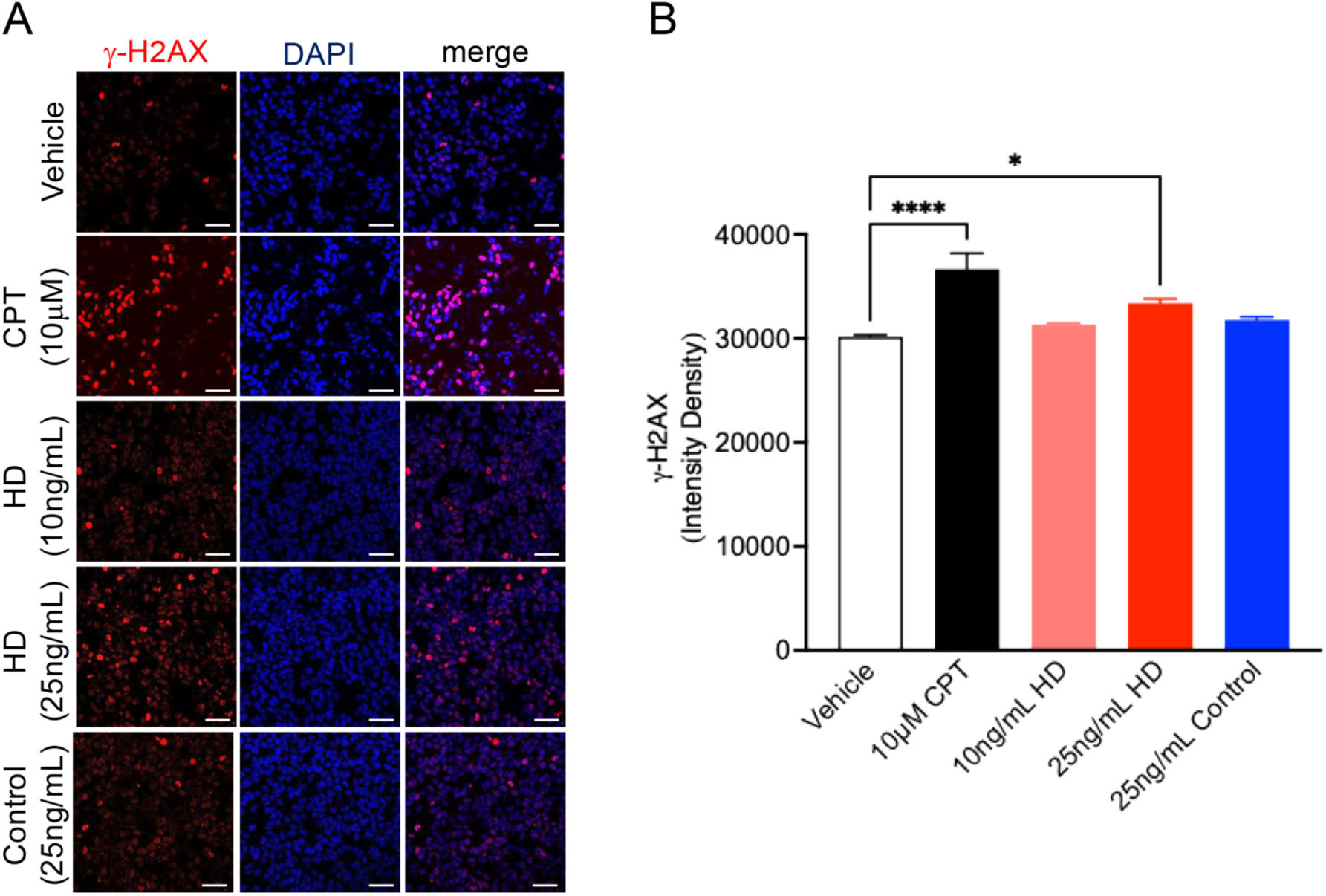
HD PFC increased γ-H2AX levels in SH-SY5Y cells. **(A)** Representative immunofluorescence images of γ-H2AX (red), DAPI (blue), and merge (right) in SH-SY5Y cells following treatment with control and HD PFC homogenates. **(B)** One-way ANOVA revealed a significant effect of treatment on γ-H2AX levels [F(4,34)=11.81; p<0.0001]. CPT treatment (10μM/2h) significantly increased γ-H2AX levels compared to vehicle-treated cells (Tukey’s test, p<0.0001). Neither treatment with 10 ng/mL of HD PFC (Tukey’s test, p=0.8218) nor treatment with 25ng/mL of control PFC (Tukey’s test, p=0.5273) altered γ-H2AX levels compared to vehicle-treated cells. In contrast, treatment with 25ng/mL of HD PFC significantly increased γ-H2AX levels (Tukey’s test, p=0.0211). Data are shown as mean±SEM. Scale bar 50μm. * p<0.05; **** p<0.0001.

**Supplementary Figure 2.**
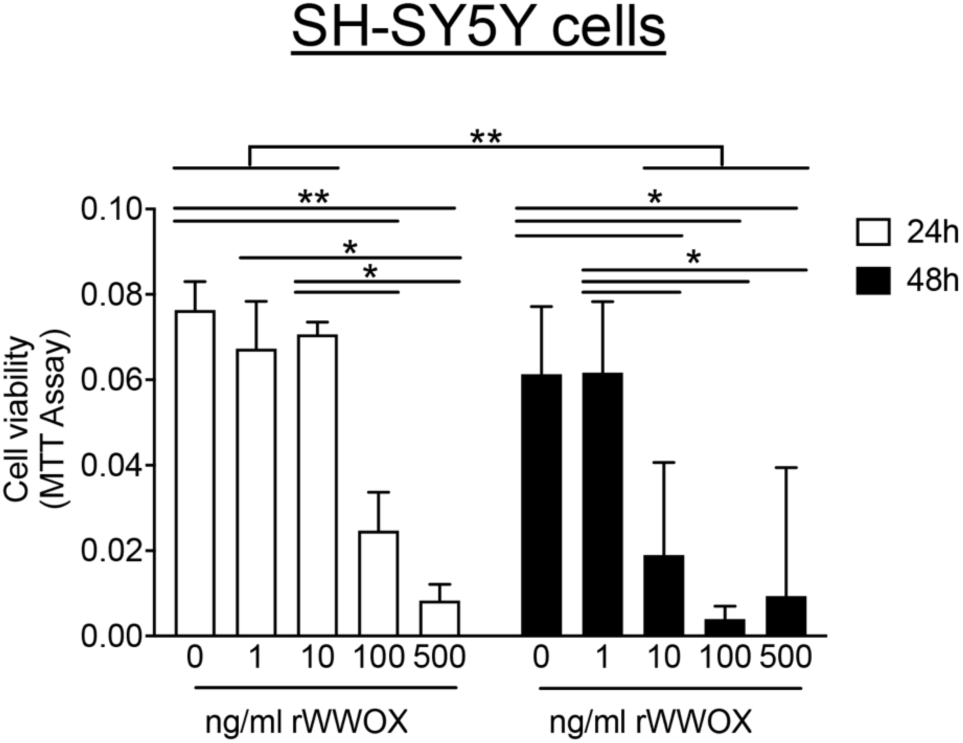
High concentrations of rWWOX reduced cell viability in SH-SY5Y cells in a dose- and time-dependent manner. SH-SY5Y cells were treated with increasing concentrations of rWWOX (1-500ng/mL) for 24h or 48h prior to measuring cell viability. Two-way ANOVA revealed a significant effect of dose [F(4,20)=21.15; p<0.0001], time [F(1,20)=10.67; p=0.0037], and dose X time interaction [F(4,20)=3.280; p=0.0320]. At 24h, treatment 100ng/mL (Tukey’s test, p=0.0086) and 500ng/mL (Tukey’s test, p=0.0004) significantly reduced cell viability compared to vehicle-treated cells, whereas 1ng/mL and 10ng/mL had no effect (Tukey’s test, p=0.9558, and p>0.9999, respectively). At 48h, treatment with 10ng/mL (Tukey’s test, p=0.0460), 100ng/mL (Tukey’s test, p=0.0030) or 500ng/mL of rWWOX (Tukey’s test, p=0.0081) significantly reduced cell viability compared to vehicle-treated cells, whereas 1ng/mL of rWWOX had no effect (Tukey’s test, p>0.9999). Data are shown as mean±SEM. Scale bar 50μm. p>0.05.

**Supplementary Figure 3.**
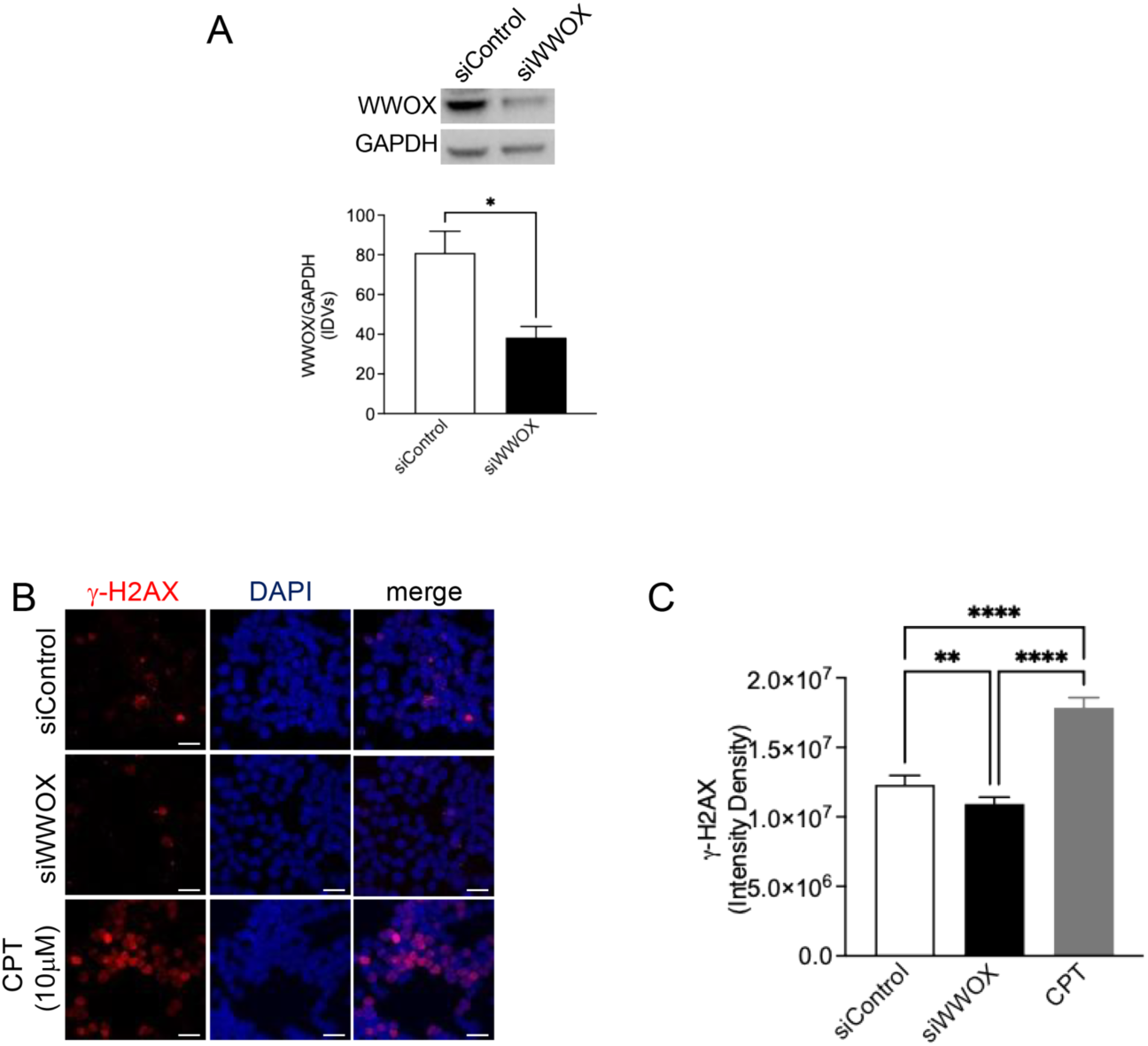
WWOX knockdown reduced γ-H2AX levels in SH-SY5Y cells. **(A)** WWOX levels were significantly decreased in siWWOX-treated compared to siControl-treated SH-SY5Y cells (n=4 replicates, Mann-Whitney U test=0, p=0.0286). **(B)** Representative immunofluorescence images of γ-H2AX (red), DAPI (blue), and merge (right) in siControl- and siWWOX-treated cells. **(C)** One-way ANOVA revealed a significant effect of treatment on γ-H2AX levels [F(2,13)=164.1; p<0.0001]. CPT treatment (10μM/2h) significantly increased γ-H2AX levels compared to both siControl- (Tukey’s test, p<0.0001) and siWWOX-treated cells (Tukey’s test, p<0.0001). In contrast, γ-H2AX levels were significantly decreased in siWWOX-treated cells compared to siControl-treated cells (Tukey’s test, p=0.0044). Data are shown as mean±SEM. Scale bar 50μm. * p<0.05; ** p<0.01; **** p<0.0001.

